# Single-cell foundation models benefit from cross-modal training: adding proteomics data beats parameter scaling

**DOI:** 10.64898/2026.08.14.744845

**Authors:** Maximilien Burq, Peter Cimermancic, Charlie Kim, Dejan Stepec

## Abstract

Leading cellular foundation models have been trained on hundreds of millions of single-cell transcriptomes, with progress increasingly driven by larger datasets and model scaling. Here, we asked whether adding a proteomics modality can improve gene-level and cell-level representations beyond scaling RNA-only models. We introduce cross-modal continued pretraining, fine-tuning a published single-cell model (Tahoe-x1) on a large corpus of proteomic profiles. Training a 70M-parameter Tahoe-x1 model for a single epoch on 48843 proteomic samples from 440 diverse mass-spectrometry studies matched or exceeded 1B- and 3B-parameter RNA-only models across most of the original Tahoe-x1 evaluation benchmarks. This shows that with the right training recipe, heterogeneous proteomics data can improve the learned representations of single-cell RNAseq samples, demonstrating strong out-of-distribution generalization. Cross-modal pretraining also improves transfer to a held-out protein perturbation benchmark, where scaling the RNA-only model does not provide comparable benefits. These results demonstrate that careful targeted curation of proteomics data can provide larger benefits than increasing the model size alone and suggest that multimodal pretraining is a promising path toward more informative biological foundation models.

## 1 Introduction

Building a “virtual cell”, a model that predicts how cells respond to genetic and chemical perturbations, is a central goal of computational biology, with direct applications to drug discovery and mechanism-of-action studies [1–3]. The dominant approach borrows from language modeling: pretrain a transformer on hundreds of millions of single-cell RNAseq profiles, and then transfer the learned representations to downstream tasks [4–6]. Recent work has focused on the model and data scale: more cells and more parameters (Tahoe-x1 to 3B; X-Cell to 4.9B), with X-Cell reporting LLM-like power-law scaling of its training loss across two orders of magnitude of model size [2].

However, increasing scale alone has not consistently translated into improved biological utility. Single-cell foundation models do not reliably outperform simple additive-linear baselines for perturbation-effect prediction [7], and their zero-shot representations can be surpassed by simpler classical approaches, such as highly variable gene (HVG) selection, principal component analysis (PCA), and scVI embeddings on integration benchmarks [8]. Moreover, recent studies suggest that pretraining performance can saturate well before the available data are exhausted, with no clear scaling relationship between the dataset size and downstream performance [9].

As parameter scaling alone has yielded diminishing improvements in biological utility, the field is increasingly exploring complementary strategies to enhance foundation models. One promising area for improving model performance is through increasing *contextual diversity* of the pretraining data (tissues, states, donors) [9, 10]. The strongest recent perturbation models advance the field by adding interventional data and richer biological priors [1, 2].

Here we introduce a modality on which such models have not previously been trained. Proteins — not transcripts—are the key molecular level at which biological functions begin to manifest. Importantly, protein abundance does not correlate perfectly with transcript levels. As a result, proteomics captures biological information that transcriptomics alone cannot, including the effects of post-transcriptional regulation. Notably, the Tahoe-x1 authors identified proteomics as a key missing modality and an important direction for future work. To our knowledge, only two studies have attempted to train models on non-structural proteomics data. ProteinTalks [11] trained a model from scratch on 18 breast cancer lines × 63 drugs (5,143 unique proteins) using proteome perturbation as the loss function. However, its performance was compared only against classical ML and never against an RNA foundation model. CAPTAIN [12] co-trained RNA and protein encoders on per-cell co-assayed CITE-seq *surface* panels (382 standardised surface proteins, 4.2M cells), with the RNA encoder initialized from scGPT. However, the scarcity of large-scale per-cell RNA – protein co-assay datasets has limited the generalizability of this approach, leaving the potential of protein representations for perturbation prediction largely unexplored.

Here, we demonstrate how proteome information can advance cell models. First, we demonstrate that crossmodal continued pretraining (CPT) using a carefully curated proteomics corpus enhances both gene- and cell-level representations within a single-cell foundation model, allowing a 70M-parameter model to outperform models up to 50× its size. Second, we establish a novel held-out protein perturbation benchmark —the first designed to evaluate single-cell foundation models beyond transcript levels—and show that incorporating proteomics data significantly improves predictions of drug-induced protein shifts. Third, through a systematic analysis of proteomics data scale and presentation strategies, we show that expanding to a complementary modality yields far greater performance improvements than simply scaling parameters in RNA-only models.

## 2. Method and results

### 2.1 Building a corpus of diverse proteomics datasets

Public proteomics datasets are not standardized for large-scale machine learning [13]. Unlike single-cell RNAseq resources, where common formats and metadata conventions have emerged, published massspectrometry proteomics studies vary widely in file structure, protein identifiers, abundance representations, and metadata availability. Sample annotations and experimental conditions are frequently embedded in manuscript text, figures, or supplementary tables rather than accompanying the deposited data. As a result, integrating public proteomics datasets requires study-specific interpretation: nearly every dataset requires custom parsing, column mapping, or metadata reconstruction (SI Table S2).

To address this challenge, we developed a curation pipeline that combines automated dataset interpretation using AI agents with expert human reviews (Figure 1A). For each dataset, agents infer file structure, identify quantitative measurements, recover sample-level metadata, and align deposited profiles with the corresponding publication. For some datasets where protein identification and quantification results were missing or computed via outdated search algorithms, we re-ran the search analysis using Tesorai Search [14]. Human experts review a random subset and ambiguous cases, including sample-condition mappings, mislabeled metadata, and experimental design details that cannot be reliably inferred from deposited files alone. The resulting pipeline enables systematic extraction, harmonization, and quality control of heterogeneous public proteomics datasets into a unified continued-pretraining corpus.

**Figure 1.**
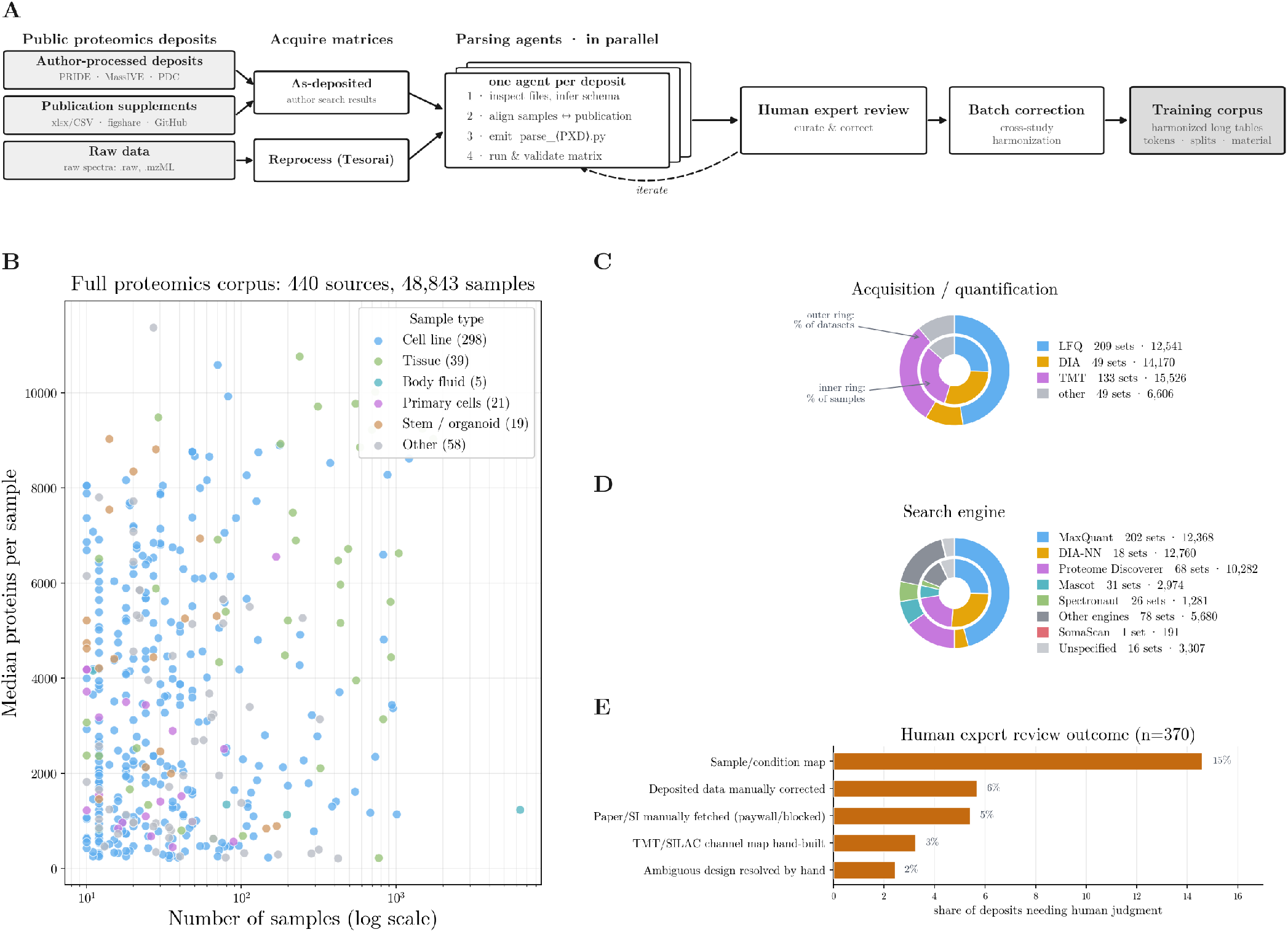
The mass-spec proteomics data used for continued pretraining (440 sources, 48,843 samples). (A) The ingestion pipeline: diverse public deposits (repository search results, publication supplements, and raw spectra) are either taken as-deposited or reprocessed through the Tesorai platform to generate quantified protein matrices. One LLM agent per deposit infers each layout and aligns samples to the source publication; human experts review and iterate with the agents; the harmonized, batch-corrected tables become the training dataset. (B) One dot per source, x = number of samples (log), y = median proteins/sample, colored by sample type. (C) The acquisition/quantification mix (LFQ / DIA / TMT / other). (D) The search engine, recovered per dataset from the PRIDE deposit metadata and source publications. In panels C and D the outer ring weights by dataset count and the inner ring by sample count. (E) Human expert review outcome: for the 370 deposits parsed by an agent, the share whose per-deposit record shows a step that needed human judgment the agent could not automate - a sample/condition mapping recovered from the paper’s prose, figures, or PDF; a manual correction of mislabeled deposited data; a paper or supplement fetched by hand; a hand-built TMT/SILAC channel map; or an ambiguous design resolved by hand. Flags are not mutually exclusive.

Using this approach, we assembled a corpus of 440 public mass-spectrometry proteomics sources comprising 48,843 samples spanning cancer types, tissues, body fluids, primary cells, organoids, and drug perturbation studies (Figure 1). Sources contributing fewer than 10 samples were excluded. The corpus captures the three dominant large-scale proteomics quantification strategies: 15,526 isobaric labeling samples (TMT/iTRAQ), 14,170 DIA samples, and 12,541 label-free samples. No single data acquisition strategy dominates the corpus, making platform heterogeneity a fundamental property of the training data.

The training data also reflects strong biological and technical coupling between sample type and acquisition strategy. Tissue samples are predominantly quantified using isobaric labeling (48%) and represent the highest complexity measurements, with a mean depth of 6,676 proteins per sample compared with 3,831 proteins per sample for cell lines. In contrast, the data from bodyfluid samples are almost exclusively DIA-based and are the lowest complexity, averaging 1,241 proteins per sample. The complete corpus composition by sample type, quantification strategy, and proteome depth is provided in SI Table S1.

### 2.2 Continued pretraining and evaluation on existing scRNAseq benchmarks

Methods We trained the Tahoe-x1 70M checkpoint, starting from their published weights and using their training recipe, for one epoch on the proteomics data. Each protein was mapped to a gene token and its abundance to the expression value. The Tahoe-x1 model’s per-sample quantile binning addressed heterogeneity in the absolute intensity scales of different proteomic datasets. We excluded drug-perturbation samples that were used as part of our evaluation datasets from the training data.

TMT data are sometimes reported as ratios between samples rather than as absolute or relative protein abundances, distinguishing them from other mass spectrometry proteomics modalities. To account for this data representation, we present each cell in both its raw and batch-corrected forms, which we refer to as a format-aware augmentation.

To test whether the improvement from proteomics continued pretraining is specific to the second modality, we built two arms that substitute or add bulk RNAseq. The RNA source is ARCHS4 [15], a uniform reprocessing of human bulk RNAseq runs from GEO and SRA. We encoded the v2.5 human atlas 877,715 of 888,821 samples (98.7%); the remainder were dropped for having <200 genes mappable to the Tahoe vocabulary. In contrast to the original Tahoe-100M dataset used for pre-training, these are bulk samples one RNAseq run per sample, from a tissue or cell line - not single cells. Each sample is tokenized to Tahoe-x1’s Ensembl gene vocabulary and utilises the same binning strategy, so a bulk sample enters the model in the same (gene, expression-bin) set format as a Tahoe-x1 cell. The RNA-only arm (“+ bulk RNAseq (1M)”) continues pretraining the 70M checkpoint on the 1,003,000 RNA samples alone for one epoch (250,750 steps at batch size 4). The combined arm (“+ proteomics + bulk RNAseq”) pools the 48,843-sample proteomics corpus with the same RNA samples and trains for one epoch over the union. Both arms match the proteomics arm’s one-epoch, per-sample budget. Source weighting is baked into the shard set at build time and read uniformly, so over a single epoch the modality mix equals the row-count ratio.

The model can be applied to either RNAseq (bulk or single-cell) or proteomics datasets to generate contextual gene-level and cell-level representations. We evaluated scRNAseq representations on three tasks: Gene essentiality prediction (DepMap [16]), Pathway membership prediction (MSigDB Hallmark [17]), and Tissue-of-origin separation (LISI [18]), using Tahoe-x1’s published evaluation code and datasets. We report PCA as a baseline comparison, and ESM2 [19] as a protein-language-model floor.

#### Results: Cross-modal CPT lifts a 70M model past the 1B and 3B

Cross-modal continued pretraining enables substantial gains in biological performance, allowing our 70M model to match or exceed the performance of the 1B and 3B checkpoints-despite being 14-fold and 50-fold smaller, respectively-across benchmarks of gene essentiality, pathway membership, and tissue-of-origin separation (Figure 2). We also evaluated continued pretraining with bulk RNAseq data, which was not included in the original Tahoe-x1 training corpus. While bulk RNAseq improved pathway membership prediction and tissue-of-origin separation, it did not improve gene essentiality prediction. Together, these findings indicate that the observed gains arise primarily from incorporating complementary, cross-modal data rather than from additional training or expanded scRNA-seq exposure.

**Figure 2.**
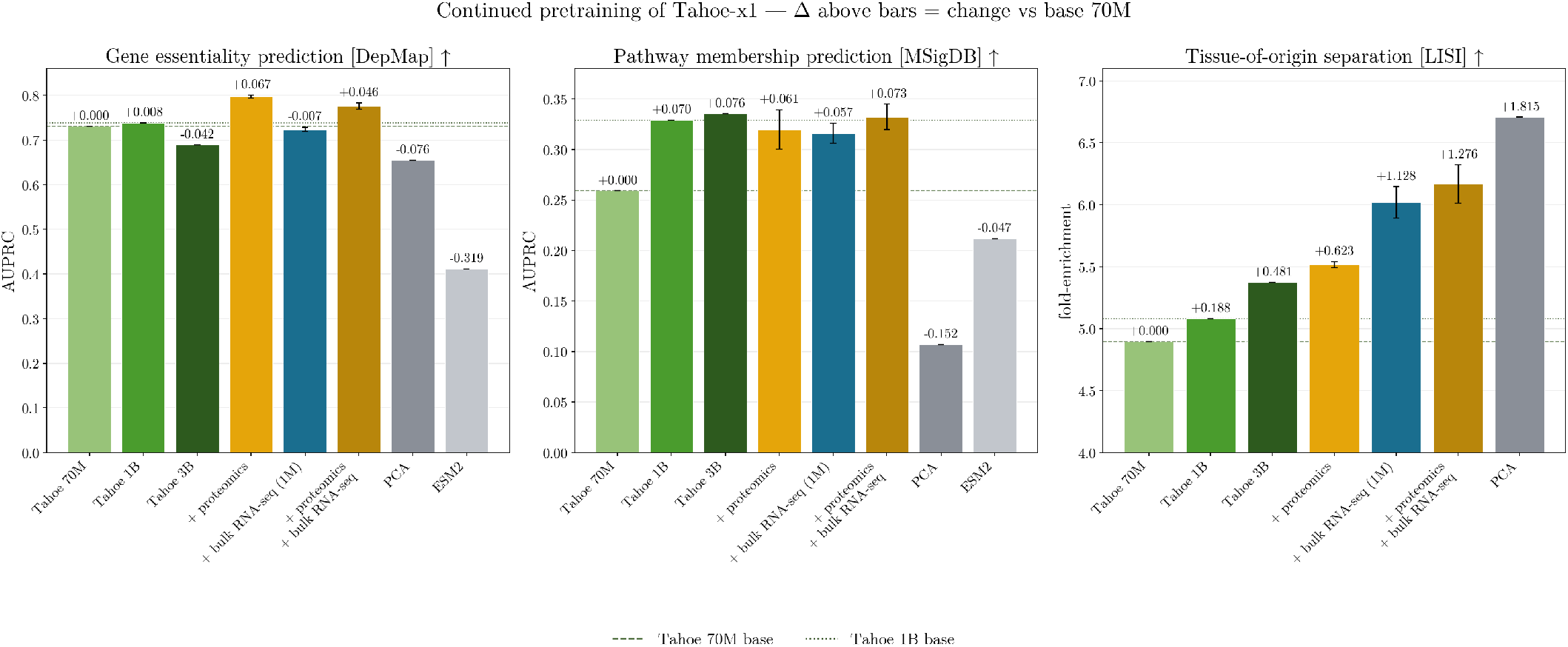
Gene essentiality prediction (DepMap AUPRC), pathway membership prediction (MSigDB AUPRC), and tissue-of-origin separation (LISI) across models: the three base Tahoe-x1 checkpoints (70M/1B/3B); 70M checkpoint with continued pretraining on proteomics (“+ proteomics”); on 1M-sample bulk RNAseq alone (“+ bulk RNAseq (1M)”); and on proteomics combined with 1M-sample bulk RNAseq (“+ proteomics + bulk RNAseq”); with PCA and ESM2 classical references. Δ above each bar = change vs base 70M; dashed/dotted lines = base 70M/1B. Error bars = *±* 1 std over 3 seeds for the multi-seed continued-pretraining arms; the base checkpoints and classical references are single runs.

A classical PCA embedding of the same data remains a strong baseline for tissue-of-origin separation, highlighting that not all representation tasks require large models. Pathway membership is the only benchmark where CPT does not improve over the larger checkpoints; it matches the 1B and 3B models within seed variability (~0.8*σ*). Even this tie demonstrates that a 70M model can reach the performance of a model 14 *×* larger with a fraction of the data and associated computational cost.

### 2.3 A held-out-drug protein-perturbation benchmark

#### Methods

To test whether cross-modal pretraining improves biological transfer beyond transcriptomic representation benchmarks, we developed a protein-level perturbation benchmark. The benchmark asks whether a model’s representation of a proteome can predict drug-induced changes in protein abundance for drugs that were not observed during pretraining. This directly evaluates whether proteomics-informed representations capture transferable biological relationships between cellular state and perturbation response.

To our knowledge, this is the first benchmark combining a single-cell foundation model, transfer to heldout drug perturbations, and a comprehensive hierarchy of trivial and classical baselines. The closest prior work, ProteinTalks, uses perturbed proteome prediction as a pretraining objective but does not evaluate heldout drug transfer. We therefore designed the benchmark to explicitly measure generalization to unseen perturbations rather than memorization of previously observed drug responses.

We construct the benchmark from two published deep drug-response proteomics datasets (Table 1). PXD014791 provides label-free (LFQ) proteomes from primary human cardiomyocytes generated from four donors and treated with 21 kinase inhibitors [20]. ProTargetMiner is a collection of TMT proteomes from three cancer cell lines (A549, MCF7, and RKO) treated with 56 anticancer drugs [21]. TMT quantifies each protein relative to a shared reference channel, so the original ProTargetMiner deposits are ratios rather than absolute abundances. Because our evaluation ranks proteins by within-sample abundance, we do not use these ratios: we instead take the absolute isotope-corrected reporter intensities from the deposited MaxQuant output and assign each channel to its drug or vehicle control using the published TMT-10 design ( [21], Supplementary Table 2), placing ProTargetMiner on the same absolute-abundance scale as the label-free cardiomyocyte proteomes.

**Table 1.** The two protein-perturbation datasets. A “pair” is one (drug, biological-context) combination. Held out pairs use drugs blocked from pretraining and scored only in evaluation (drugs listed in SI). *Pairs are reported as total / held-out.

|  | PXD014791 | ProTargetMiner |
| --- | --- | --- |
| Biological context | primary human cardiomyocytes | Cancer cell lines |
| Quantification | LFQ | TMT (absolute reporter) |
| Contexts | 4 donors | 3 cell lines (A549, MCF7, RKO) |
| Drugs | 21 | 56 |
| Samples | 237 drug / 71 control | 237 drug / 35 control |
| Median depth | 2,780 proteins | 3,746 proteins |
| Pairs* | 60 / 58 | 73 / 61 |

#### Protocol

We evaluate transfer using leave-one-pair-out cross-validation over drug-context pairs within each dataset. For each held-out pair, we embed all available proteomic samples, aggregate embeddings within each pair, and predict the perturbed proteome using the cosine-similarity-weighted mean response of the k=5 nearest training pairs in embedding space. Each prediction is evaluated relative to the held-out pair’s matched control samples (vehicle or DMSO-treated samples from the same biological context), ensuring that the benchmark measures prediction of the drug-induced proteomic response rather than similarity in baseline proteome composition. To assess transfer beyond the exact perturbation contexts encountered during pretraining, we partition drug-context pairs into “seen” and “held-out” groups based on whether that specific combination was present in the continued-pretraining corpus, and report results on held-out pairs throughout.

#### Metrics

For each held-out pair, we compare the predicted proteomic response with the measured response relative to the matched control. Given predicted proteome *p*, measured proteome *r*, and matched control *c*, we define the predicted and measured log-fold-changes as 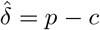 and *δ* = *r −c*.

Because mass-spectrometry proteomics measures only a subset of the proteome (typically *~*5-10k proteins per sample), we restrict evaluation to proteins detected (*>* 0) in either the predicted or measured response. Including proteins absent from both profiles would artificially inflate agreement by rewarding shared missingness rather than biological prediction.

We report two correlation metrics:

- **Full-proteome Pearson** 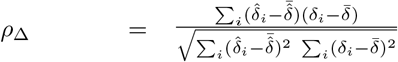, across all detected proteins.
- **Top-100 DE Pearson:** Pearson correlation restricted to the 100 proteins with the largest measured absolute response |*δ*_*i*_|. This metric emphasizes proteins most affected by the perturbation, since most proteins remain relatively unchanged and whole-proteome averages can dilute perturbation-specific signals.

Whole-proteome metrics can therefore favor predictors that capture broad expression structure while failing to identify the proteins that respond to perturbation. To reduce this signal dilution, we use top-100 DE Pearson as our primary metric and report full-proteome Pearson as a robustness analysis. Both metrics are averaged across held-out drug-context pairs.

#### Baselines

Because the target is a *perturbation effect* rather than absolute proteomic states, we include several kinds of baseline: one that predicts no change, one that ignores the embedding and predicts the mean drug effect, and classical embeddings run through the same predictor as the models. Mejia et al. [22] argue that perturbation benchmarks usually lack exactly this calibration. Without it, a metric cannot tell a model that has learned perturbation-specific structure from one that has collapsed to the dataset mean. For held-out pair *i* (drug *d*_*i*_, context *g*_*i*_, matched control *c*_*i*_), let *δ*_*j*_ = *r*_*j*_ *− c*_*j*_ be the measured log-fold-change of any other pair *j*.

- **control-mean:** predict no change, 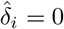 (i.e. *p*_*i*_ = *c*_*i*_). Its per-pair Pearson is undefined (zero variance) and reports as NaN; it fixes the floor.
- **perturbation-mean:** ignore the embedding and predict the average perturbed proteome over all other pairs, 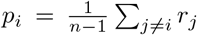. Tests whether the_*ij*_ embedding beats “predict the mean drug effect.”
- **HVG:** the same cosine-kNN predictor as the models (*k* = 5), but the embedding is the 2,000 highest-variance proteins of the proteome (L2-normalised).
- **PCA:** the same cosine-kNN predictor, with a 512-component PCA of the log-proteome as the embedding.
- **additive-linear** [7]: predict the effect as a two-way additive decomposition, 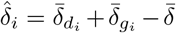, where 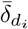 and 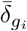 average the measured *δ* over the other pairs sharing *i*’s drug and its context, and 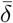 is the global mean; 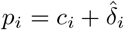.

The control-mean, perturbation-mean, and additive-linear baselines do not use learned representations. HVG and PCA use the same kNN prediction procedure as the foundation models, isolating the contribution of the learned embedding itself.

#### Cross-modal CPT improves protein transfer; RNA scale does not

To evaluate whether cross-modal pretraining improves transfer at the protein level, we scored every model against the baselines (Figure 3). On the primary top-100 DE metric, proteomics CPT improves the performance (measured by correlation) of the 70M model by +0.070 on the cardiomyocyte dataset (0.542 *→* 0.611) and by +0.067 on the cancer-cell-line dataset (0.746 *→* 0.813). In contrast, increasing RNA-only model scale does not provide comparable gains: the 1B base model underperforms the 70M base model on both datasets (0.515 and 0.718, respectively). The full-proteome metric shows the same trend, with slightly larger improvements from CPT with proteomics data (+0.083 and +0.088). Thus, the transfer benefit is associated with protein-level continued pretraining rather than increased RNA model scale. The weak performance of baselines on ProTargetMiner reflects its heterogeneity, which spans three distinct cancer cell lines: predicting an average response poorly captures context-specific perturbation effects. As a result, absolute performance on this dataset reflects both biological-context representation and drug-response prediction.

**Figure 3.**
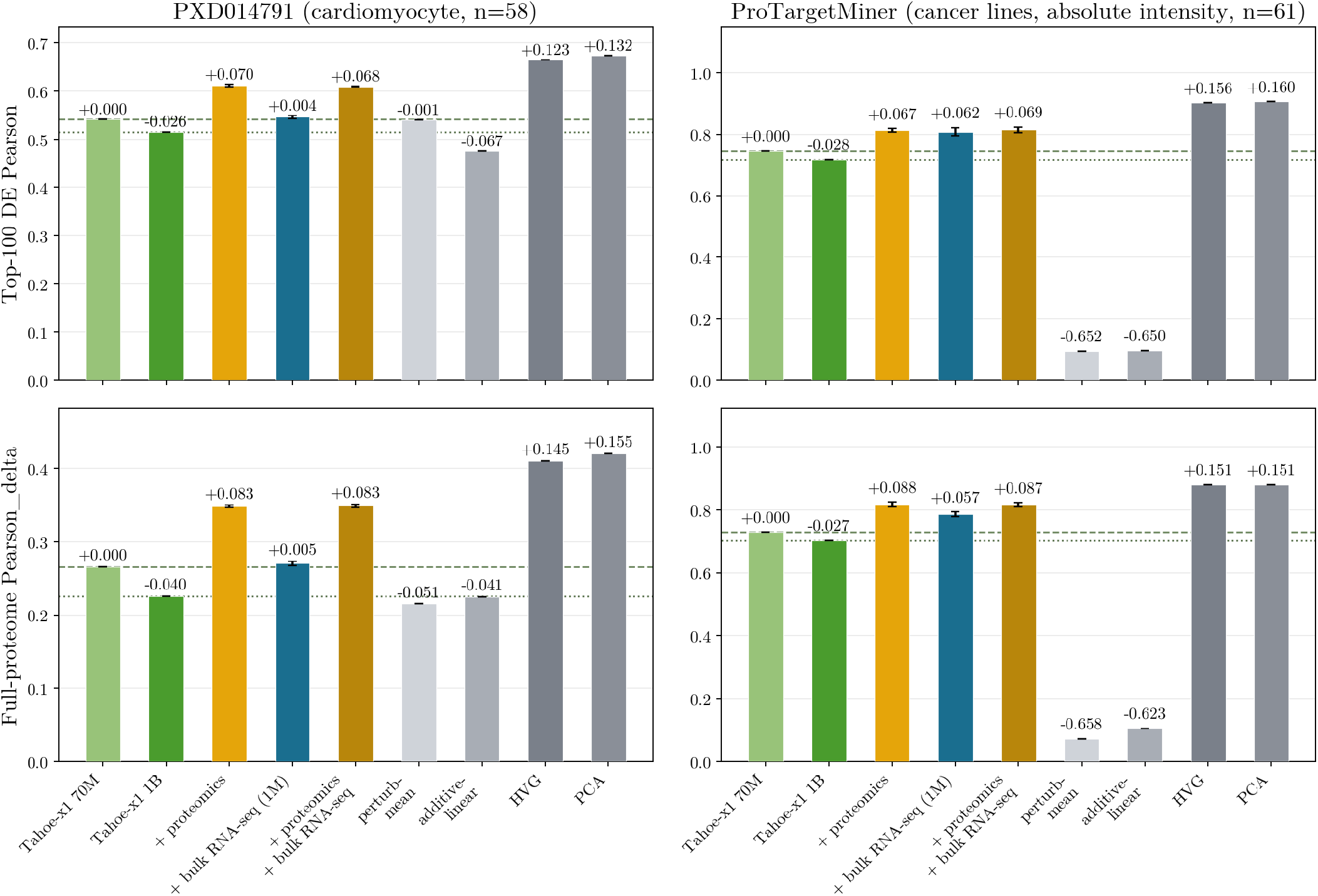
Protein-perturbation transfer on cardiomyocyte-donor (PXD014791) and cancer cell line (ProTargetMiner) proteomes, for the base 70M and 1B, the continued-pretraining arms (proteomics; 1M-sample bulk RNAseq alone; and proteomics combined with 1M-sample bulk RNAseq), and the perturbation baselines (grey): perturbation-mean, the additive-linear model, HVG, and PCA. Leave-one-pair-out correlation of predicted vs measured per-protein log-fold-changes; top row = top-100 DE proteins (the primary metric), bottom row = full proteome. The control-mean baseline (predict no change) is undefined under this correlation metric and is not shown. ProTargetMiner is scored against absolute reporter intensities. Held-out drug pairs are identical across models: PXD014791 n=58, ProTargetMiner n=61. Δ above bars = change vs base 70M; dashed/dotted = base 70M/1B. Error bars = *±*1 std over 3 seeds for the continued-pretraining arms; the base checkpoints and single-run baselines carry none.

### 2.4 Protein transfer improves with corpus size

To assess the effect of biological data scale, we varied the amount of proteomic information available during continued pretraining (Figure 4). We trained on random subsamples ranging from 300 to all 48,843 samples, using uniform sampling and a single epoch-the same configuration as the full CPT proteomics model-with three seeds per condition. Gene essentiality improved from 0.751 *±* 0.005 at 300 samples to 0.796 *±* 0.002 on the full corpus, exceeding the 1B RNA-only base-line (0.739) from the smallest subsample tested. Tissue-of-origin separation surpassed the 1B baseline (5.08) at approximately 5,000 samples. Pathway membership matched the 1B checkpoint (0.329) at any 25,000 samples, consistent with this being the benchmark where cross-modal CPT provides the smallest improvement. Both protein perturbation benchmarks showed increasing performance as we increased the amount of available data, suggesting that additional data curation efforts to scale the available proteomics data could yield further improvements.

**Figure 4.**
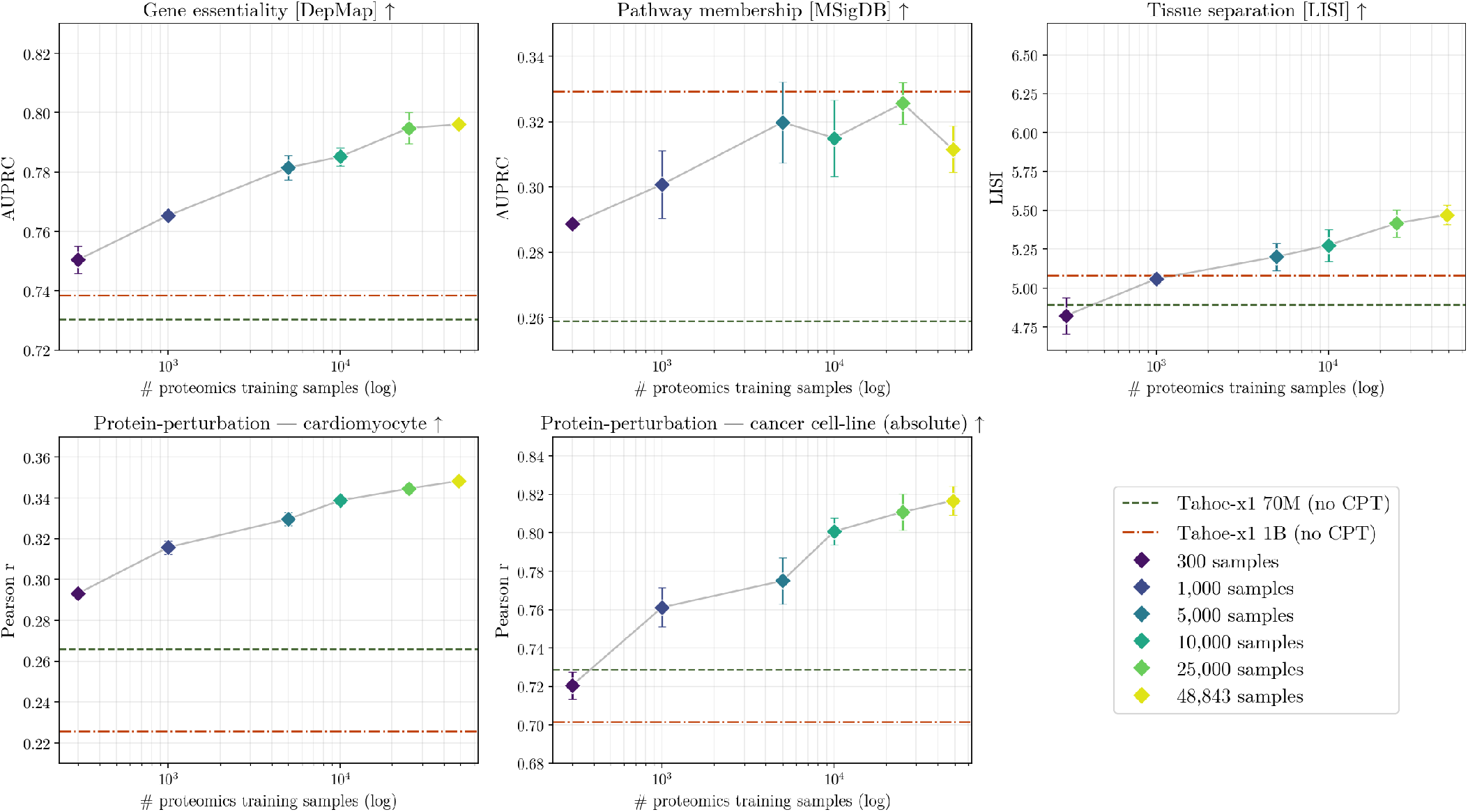
Performance vs number of proteomics continued-pretraining samples (log scale; random subsamples of the corpus, trained for one epoch), for gene essentiality (DepMap AUPRC), pathway membership (MSigDB AUPRC), tissue separation (LISI), and protein-perturbation transfer (cardiomyocyte, and cancer-cell-line scored on absolute intensities). Three seeds at every scale; error bars = *±*1 std over seeds. Dashed lines = base Tahoe-x1 70M and 1B.

## Data and code availability

The evaluation code for the protein-perturbation benchmark - the curated proteomes, the leave-one-pair-out splits, the baseline implementations, and the scoring code - is available at https://github.com/tesorai/cross_modal_foundation_model. The continued-pretrained model weights are available at https://huggingface.co/Tesorai/Trove1-70m.

## Supplementary Information

**Table S1.** Corpus composition by sample type: number of datasets, samples, sample-weighted mean depth (proteins/sample), and the quantification mix in samples (the four mix columns sum to the sample count). Sample type is assigned per dataset from the observed sample labels; the quantification family is a coarse grouping of ~110 free-text labels, so it reflects the recorded quantification type rather than an instrument-level acquisition annotation. other (6,606 samples, 13.5%) is left unassigned rather than guessed at, and comprises ratio or pre-normalised values that cannot be attributed to a family (3,114), SILAC and other metabolic labelling (2,566), sources with no quantification type recorded (565), and assay-specific readouts such as kinobeads, BioID, phospho-site sums and one aptamer panel (361). Unclear is a sample type we could not determine from the labels: sources mixing patient and cell-line samples, and sources whose sample label is empty.

| Sample type | Datasets | Samples | % of samples | Mean depth | LFQ | DIA | TMT | other |
| --- | --- | --- | --- | --- | --- | --- | --- | --- |
| Cell line | 298 | 21,910 | 44.9% | 3,831 | 9,833 | 2,870 | 7,059 | 2,148 |
| Tissue | 39 | 15,023 | 30.8% | 6,676 | 338 | 3,522 | 7,188 | 3,975 |
| Body fluid | 5 | 6,578 | 13.5% | 1,241 | 12 | 6,555 | 11 | – |
| Primary cells | 21 | 723 | 1.5% | 2,856 | 377 | – | 139 | 207 |
| Stem / organoid | 19 | 741 | 1.5% | 3,350 | 541 | 30 | 158 | 12 |
| Unclear | 58 | 3,868 | 7.9% | 2,406 | 1,440 | 1,193 | 971 | 264 |
| **All** | **440** | **48,843** | **100.0%** | **4,223** | **12,541** | **14,170** | **15,526** | **6,606** |

## Data-parsing complexity (SI TableS2)

Because published proteomics deposits share only high level conventions for formatting, we developed a dedicated parser for each deposit type. We scored the ingest requirements of every dataset along seven binary dimensions. The seven dimensions were not rare edge cases: the curated datasets triggered a median of 4 (mean 4.1) of the 7, 99% trigger at least one, and only 5 of 439 were trivially parseable (all seven FALSE).

**Table S2.**
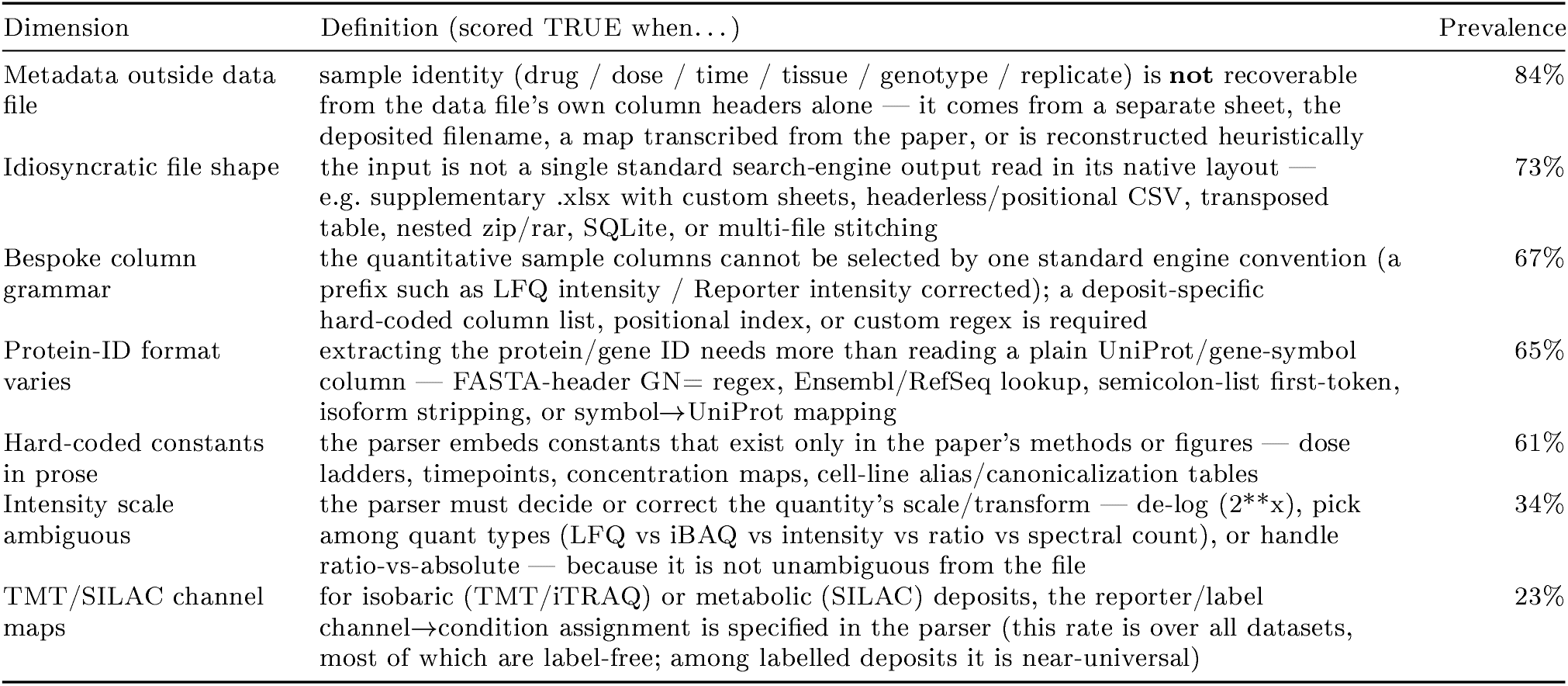
The seven data-parsing complexity dimensions: definition and measured prevalence across the 439 scored datasets.

### Proteomics perturbation benchmark

PXD014791 — 21 of 22 drugs held out: Afatinib, Axitinib, Bosutinib, Cabozantinib, Dabrafenib, Dasatinib, Erlotinib, Gefitinib, Imatinib, Lapatinib, Nilotinib, Pazopanib, Ponatinib, Regorafenib, Ruxolitinib, Sorafenib, Sunitinib, Tofacitinib, Trametinib, Vandetanib, Vemurafenib.

ProTargetMiner — 47 of 56 drugs held out: 2-methoxyestradiol, 5-fluorouracil, Afatinib, Apatinib, Auranofin, Axitinib, Azacitidine, Bortezomib, Bosutinib, Cabozantinib, Camptothecin, Carmofur, Crizotinib, Dasatinib, Docetaxel, Doxorubicin, Enzalutamide, Epirubicin, Etoposide, Everolimus, Floxuridine, Fludarabine, Gefitinib, Genistein, Idarubicin, Irinotecan, Lapatinib, Methotrexate, Mitotane, Nilotinib, OSI-420, Oxaliplatin, Paclitaxel, Pazopanib, Ponatinib, RITA, Raltitrexed, Regorafenib, Ruxolitinib, Sorafenib, Sunitinib, Temsirolimus, Teniposide, Topotecan, Vemurafenib, Vincristine, Vismodegib.

Drugs kept in training (not scored)

- PXD014791 (1): Trastuzumab.
- ProTargetMiner (9): 8-azaguanine, Azaguanine, Bleomycin, Lomustine, Nutlin, OSW-1, Tri-1, Tri-2, b-AP15.

### The benefits of proteomics are robust to training dataset composition

Given the heterogeneity of our curated proteomics dataset, one natural question is to ask whether some sample types and preprocessing steps contribute more than others to the increase in performance. To answer this, we fixed the corpus and one-epoch training budget and varied individual data type weighting choices relative to the full “+ proteomics” corpus (format-aware augmentation with uniform sampling). Across all tested variants, continued pretraining remained substantially better than both RNA-only base checkpoints on every benchmark (FigureS1).

Removing the batch-corrected view and training only on the raw proteomic representation produced a small but consistent decrease on gene-level benchmarks: gene essentiality decreases from 0.797 to 079l and pathway membership from 0.320 to 0.316. Thus, presenting both raw and corrected views provides a modest but reproducible benefit compared with either representation alone.

We also tested a sampling strategy where we over-sample under-represented cell-lines (1,800 total) to increase the diversity of contexts seen during training. It increases gene essentiality from 0.797 to 0.808 and tissue separation from 5.49 to 6.04, compared with 4.89 for the base 70M model, while leaving pathway membership and perturbation benchmarks largely unchanged. We retained the simpler uniform-sampling configuration as the headline model to maintain consistency with the arm combining proteomics and bulk RNAseq. In contrast, directly oversampling individual material classes (tissue, cell line, or body fluid) by 2*×* did not produce measurable improvements across benchmarks.

**Figure S1.**
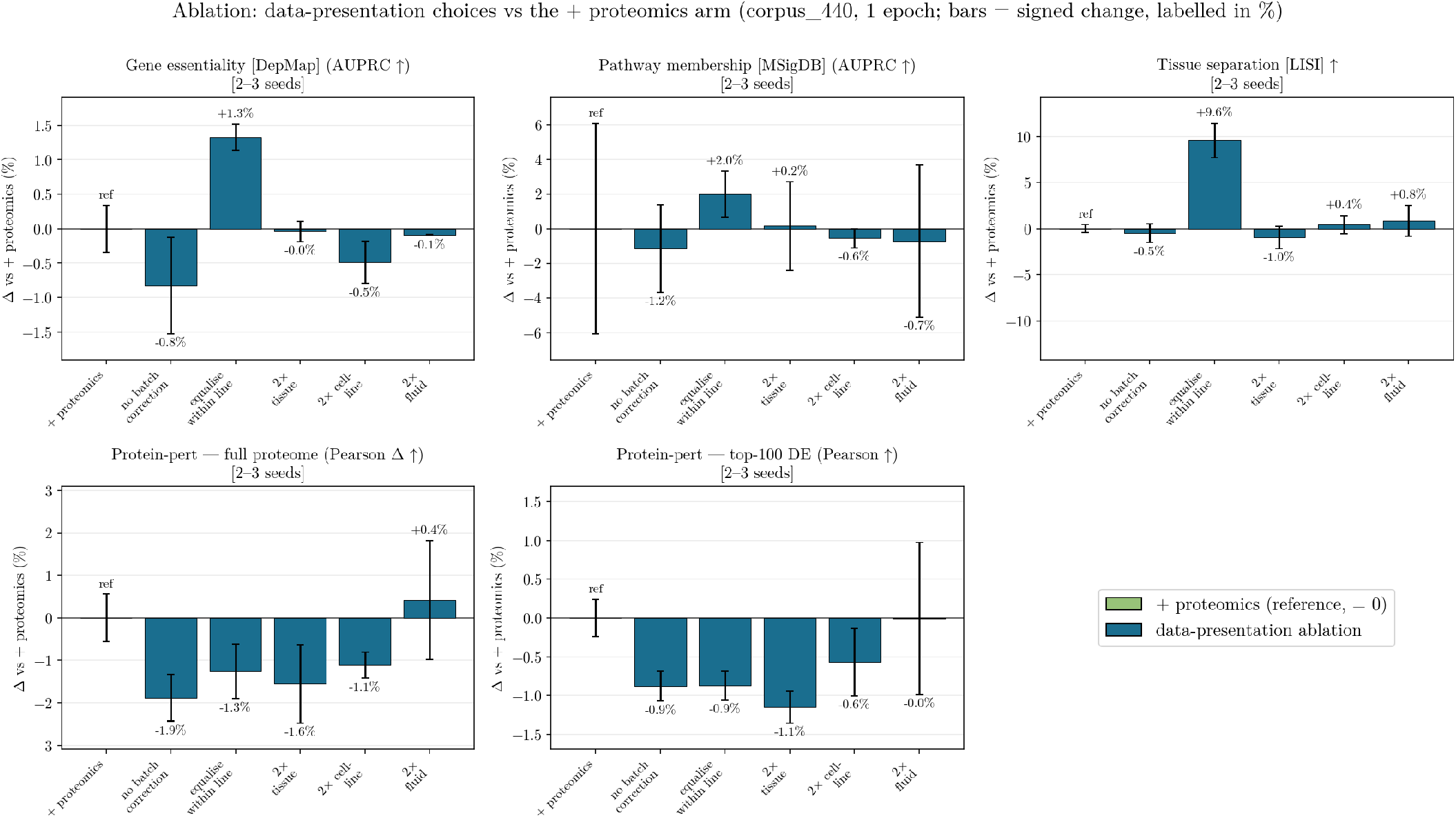
Gene essentiality (DepMap AUPRC), pathway membership (MSigDB AUPRC), tissue separation (LISI), and protein-perturbation transfer on the cardiomyocyte proteome (full-proteome and top-100 DE), for the headline + proteomics arm and a set of data-presentation ablations around it, all on the full corpus (one epoch). Bars: hatched grey = reference models (base 70M/1B); green = the + proteomics headline arm (format-aware augmentation, uniform sampling); blue = data-presentation ablations (no batch-correction view; equalise-within-line sampling; and 2*×* oversampling of tissue / cell-line / body-fluid samples). Error bars denote *±*1 std over seeds (3 for the gene-embedding panels; 2-3 for protein-perturbation, per panel); the reference bars are deterministic single runs.

## Notes

https://github.com/tesorai/cross_modal_foundation_model

https://huggingface.co/mburq/Trove-1-70m

### Competing Interest Statement

Authors are employees and Shareholders of Tesorai, Inc

